# Chromosome-level genome assembly and annotation of the social amoeba *Dictyostelium firmibasis*

**DOI:** 10.1101/2024.02.22.581529

**Authors:** Bart Edelbroek, Jonas Kjellin, Jon Jerlström-Hultqvist, Sanna Koskiniemi, Fredrik Söderbom

**Affiliations:** Department of Cell and Molecular Biology, BMC, Uppsala University, SE-751 24, Uppsala, Sweden

## Abstract

*Dicytostelium firmibasis* is a member of Dictyostelia, a group of social amoebae that upon starvation display aggregative multicellularity where the amoebae transition from uni-to multicellular life. The *D. firmibasis* genome assembly that is currently available is of limited use due to its low contiguity, large number of undetermined bases, and lack of annotations. Here we used Nanopore long read sequencing, complemented with Illumina sequencing, and developmental transcriptomics as well as small RNA-sequencing, to present a new, fully annotated, chromosome-level *D. firmibasis* genome assembly. The new assembly contains no undetermined bases, and consists mainly of six large contigs representing the chromosomes, as well as a complete mitochondrial genome. This new genome assembly will be a valuable tool, allowing comprehensive comparison to *Dictyostelium discoideum*, the dictyostelid genetically tractable model. Further, the new genome will be important for studies of evolutionary processes governing the transition from unicellular to multicellular organisms and will aid in the sequencing and annotation of other dictyostelids genomes, many of which are currently of poor quality.

## Background & Summary

*Dictyostelium firmibasis* is a member of Dictyostelia, a phylogenetic group of dictyostelid social amoebae^1^. These organisms are unicellular and free-living while food is plentiful, but at the onset of starvation, about 100,000 amoebae can stream together to start a multicellular developmental program. The cells differentiate and go through distinct morphological stages, culminating in a fruiting body where 20% of the cells sacrifice themselves to form a stalk that elevates the remaining cells, which sporulate. The spores are resistant to environmental stress and await dispersal to more favorable places^2–4^.

Dictyostelids have proven to be excellent models due to their peculiar lifecycle, providing insight into the evolution of multicellularity and altruism^2,5^. Furthermore, one of the dictyostelids, the genetically tractable organism *Dictyostelium discoideum*, is used as a model for bacterial infections and to uncover the molecular mechanisms behind biological processes such as chemotaxis and phagocytosis ^6–9^. In addition, the function and evolution of non-coding (nc)RNAs, such as micro(mi)RNAs, have been studied in Dictyostelia and is a main focus of our research^10–15^. Since small RNAs, such as miRNAs, are short (commonly 21 nt) and at least in *D. discoideum*, derived from AT-rich, hard to sequence intergenic regions, high quality complete genomes are essential for ncRNA studies. *Dictyostelium firmibasis* is of particular interest because it is closely related to *D. discoideum*, which has been extensively studied over the last century and was among the first protists with a fully sequenced and annotated genome^4,16^. A rough *D. firmibasis* genome assembly was available already 2012^17^, and has been valuable for comparison with *D. discoideum* or other dictyostelids to study evolution^10,18–21^. The assembly however lacks annotations, is fragmented, and contains many unresolved gaps, which limits to what extent *D. firmibasis* can be studied. Instead, a high-qualitative chromosome-level reference genome with gene annotations would give a more complete understanding of this organism and enable comprehensive comparison to other dictyostelids.

One of the challenges with sequencing *D. firmibasis, D. discoideum* or other closely related social amoebae is that they are particularly AT-rich, especially in intergenic regions^18,22^. This is illustrated by the *D. discoideum* genome, which has an AT-content of 86.2% in intergenic regions^19^. Not only do these repetitive stretches of adenine and thymine hinder resolving the bases during sequencing, it also complicates assembly^16^. However, advances in long-read sequencing (third-generation sequencing) have made it feasible to get high coverage of the genome, including intergenic regions, to allow for a more complete assembly^23,24^.

In this study, we sequenced the genome of the *D. firmibasis* TNS-C-14 strain^25^. We acquired 7.5 Gbp of Oxford Nanopore long-read sequences and 24.9 Gbp of Illumina short-read sequences and were able to de-novo assemble the *D. firmibasis* genome. This resulted in six main contigs of 9.4 Mbp to 3.9 Mbp and six small contigs of 118 kbp to 24 kbp, for a combined assembly size of 31.5 Mbp with 599 times total coverage (Fig. 1). Analysis of the telomeres and comparison to the *D. discoideum* genome confirmed that the six large contigs likely represent the complete chromosomes. The remaining contigs include the fully assembled mitochondrial DNA, the linear extrachromosomal DNA harboring ribosomal DNA, and DNA belonging to the *Dictyostelium* Intermediate Repeat Sequence 1 (DIRS1) retrotransposon. By performing transcriptomics at three different stages during the lifecycle of *D. firmibasis*, we could annotate 11044 genes (Fig. 1). For 92% of these, homologs in *D. discoideum* were identified, and in most cases their expression matched that of *D. firmibasis* during development.

**Fig. 1.**
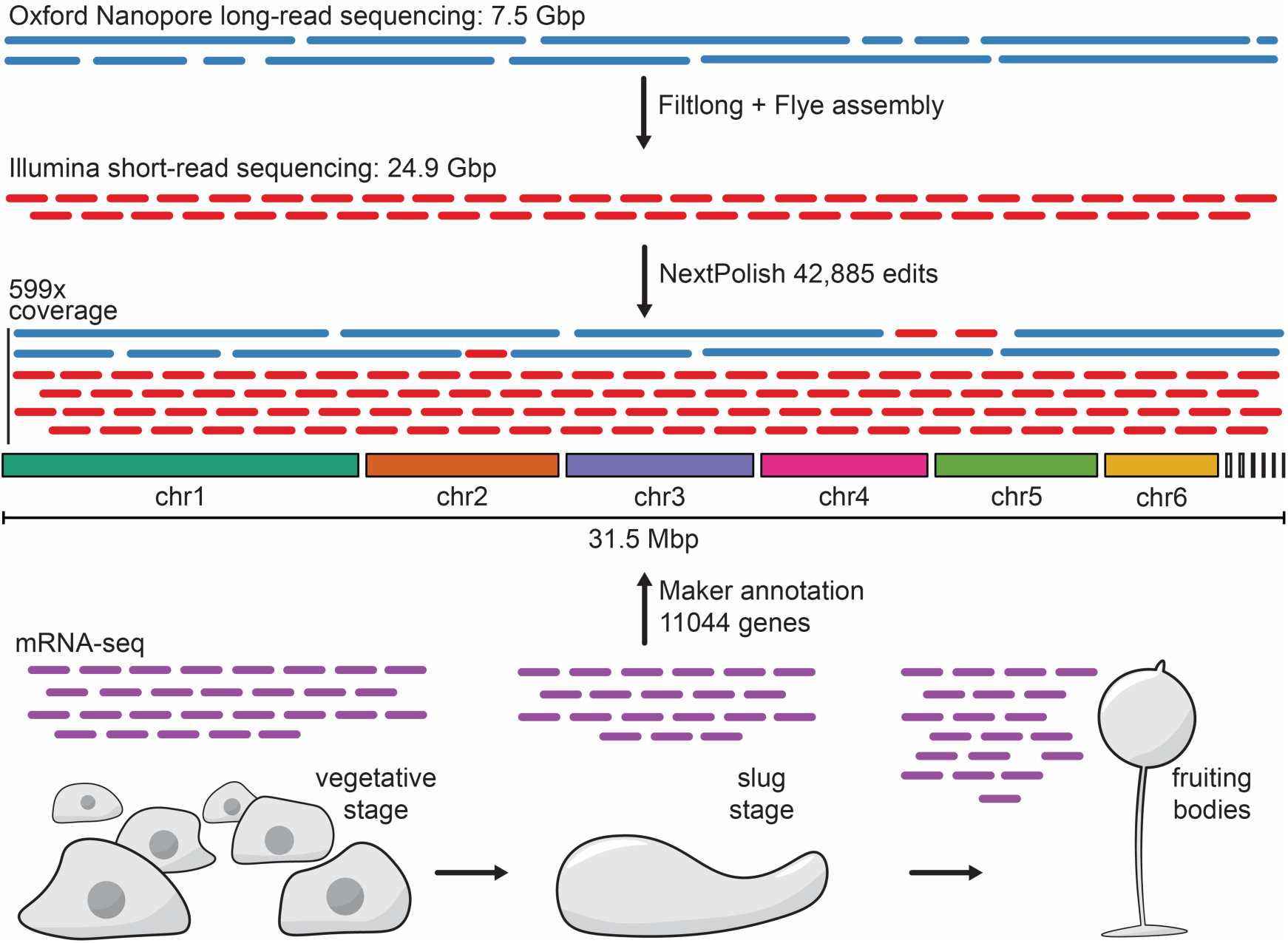
Overview of *D. firmibasis* whole genome sequencing and annotation project. De-novo assembly of *D. firmibasis* genome from 599x long- and short-read coverage resulted in 31.5 Mbp chromosome-level genome assembly, consisting of 12 contigs. mRNA-seq was performed from three distinct morphological stages (vegetative stage, slug stage, fruiting bodies) to capture also temporally expressed transcripts. For details, see Methods.

## Methods

### Amoebae single cell selection and growth

*D. firmibasis* TNS-C-14 was obtained from the Dicty Stock Centre^25^ (DSC; Strain ID DBS0235812). Approximately 20 cells were scraped from a plaque and resuspended in 250 μl *Escherichia coli* 281 culture grown at 200 rpm, 37°C, O/N (DSC Strain ID DBS0305927). The bacteria-amoeba mix was plated on 96 mm SM Agar/5 (Formedium) and grown at 22°C for 3 days. *D. firmibasis* cells were collected from the rim of the plaques formed and were verified by amplifying and sequencing the 18S ribosomal RNA (rRNA) gene with primers 5’-GTTTGGCCTACCATGGTTGTAA-3’ and 5’-CACCTCTCGCCCCAATATGA-3’. Genomic DNA, used as template, was isolated as described previously^26^ but with Triton-X100 (Sigma-Aldrich) instead of NP40. The sequenced PCR-product was aligned to the *D. firmibasis* TNS-C-14 18S rRNA gene reference (GenBank: AM168041.1).

For genomic DNA isolation for long- and short-read sequencing, approximately 10^5^ cells originating from a validated plaque, were resuspended in 750 μl *E. coli* 281 O/N culture as described above, plated on three SM Agar/5 plates and grown for 30 h until the majority of bacteria had been consumed and *D. firmibasis* started streaming together to form aggregates. Amoebae from the three plates were harvested using Nunc Cell Scrapers (Thermo Fisher) into 50 ml PDF buffer (20 mM KCl, 9.2 mM K_2_HPO4, 13 mM KH_2_PO_4_, 1 mM CaCl_2_, 2.5 mM MgSO_4_). Cells were harvested at 400xg for 5 min, and washed five times with 50 ml PDF buffer to reduce the number of bacteria.

For mRNA-seq, both *D. firmibasis* and the *D. discoideum* AX2-RRK strain (DSC Strain ID DBS0235521) were grown as above on three SM Agar/5 plates with *E. coli* 281, however after harvesting and washing, one third of the vegetative stage cells were resuspended in 1 ml TRIzol Reagent (Invitrogen) and stored at −20°C until further processing. One third of the cells was plated onto NN-Agar [8.8 mM KH_2_PO_4_, 2.7 mM Na_2_HPO_4_, 15 g/L Agar] and developed until the slug stage. The remaining third of the cells was plated onto NN-Agar and developed until the fruiting body stage. The developed cells were harvested using Nunc Cell Scrapers, resuspended into 1 ml TRIzol Reagent and stored at −20°C until further processing. Three biological replicates were prepared for each stage, for a total of 9 mRNA-seq libraries per strain.

### Genomic DNA preparation and sequencing

Genomic DNA was isolated using the Genomic-tip 100/G columns (Qiagen) according to “Protocol: Preparation of Cell Culture Samples” from the Qiagen Genomic DNA Handbook, with the exception of 0.2 mg/ml RNase A (Thermo Scientific) added during lysis of the nuclei in the supplied G2 buffer. 6*10^8^ cells were used as input, which yielded 18.6 μg genomic DNA. For long-read sequencing, an initial clean-up was performed by adding 250 μl of *D. firmibasis* genomic DNA (9.3 μg) to 112.5 μl AMPure XP Beads (Beckman Coulter) giving a ratio of 1:0.45 (V/V), to select for high molecular weight DNA. The DNA was bound to the magnetic beads by rotating end-over-end for 5 min, washed twice with 200 μl 70% EtOH and eluted in 62 μl H_2_O by incubating at 37°C for 10 min, after which the beads were magnetically removed. Further selection of high molecular weight DNA was performed using the Short Read Eliminator (SRE) kit (PacBio) according to the manufacturer’s guide, which yielded 3.2 μg of high molecular weight DNA. Long-read sequencing libraries were prepared from 1.5 μg genomic DNA with the SQK-LSK112 Ligation Sequencing Kit (Oxford Nanopore) according to manufacturer’s protocol. 100 ng of the resulting library was sequenced on an R10.4 flowcell (Oxford Nanopore) on the MinION Mk1C (Oxford Nanopore) for 72 h. Basecalling was performed with Guppy v6.3.2 (Oxford Nanopore) using the r104_e81_sup_g610 model. In total, 7.5 Gbp of long-read sequences were generated (Fig. 1).

For short-read sequencing, 4.5 μg *D. firmibasis* genomic DNA was cleaned with AMPure XP Beads as described above but at a 1:0.5 (V/V) ratio. The sequencing library was prepared with 1 μg of DNA using the TruSeq PCR-free DNA sample preparation kit (cat# 20015962, Illumina), according to the TruSeq DNA PCR-free Reference Guide, using unique dual indexes (cat# 20022370, Illumina) and targeting an insert size of 350 bp. The library was sequenced on a NovaSeq 6000 system (Illumina), with paired-end 150 bp read lengths, on an SP flowcell using v1.5 sequencing chemistry, which resulted in 24.9 Gbp of short-read sequences (Fig. 1). Library preparation and short-read sequencing were performed at SciLifeLab Uppsala.

### mRNA preparation and sequencing

RNA was isolated from three developmental stages of *D. firmibasis* and *D. discoideum*, using TRIzol Reagent (Invitrogen), according to the User Guide, with an additional 75% EtOH wash of the RNA pellet. 4 μg of RNA was DNase treated using DNase I, RNase-free (Thermo Scientific) according to manufacturer’s protocol. The RNA was purified by mixing equal volumes of RNA and Phenol:Chlorofrom:Isoamyl alcohol 25:24:1 (PanReac AppliChem) by vigorous shaking for 15 s, incubating 3 min at 21°C, followed by centrifuging at 12,000xg for 15 min at 4°C. The upper aqueous phase was added to three volumes 99% EtOH, 0.1 volumes 3M NaOAc, incubated on ice for 90 min and centrifuged at 12,000xg for 30 min at 4°C. EtOH was discarded and the RNA pellet was washed with five volumes 70% EtOH with centrifugation at 12,000xg for 15 min at 4°C whereafter the pellet was airdried for 2 min, and dissolved in H_2_O.

mRNA sequencing libraries were prepared from 500 ng total RNA using the TruSeq stranded mRNA library preparation kit (cat# 20020595, Illumina) according to manufacturer’s protocol, including polyA selection. Unique dual indexes (cat# 20022371, Illumina) were used. The libraries were sequenced on a NovaSeq 6000 system (Illumina), with paired-end 150 bp read lengths, on an SP flowcell using v1.5 sequencing chemistry. mRNA library preparation and sequencing were performed at SciLifeLab Uppsala.

### Genome assembly and polishing

Adapter trimming and read-splitting of long-reads was performed using Guppy v6.3.2 (Oxford Nanopore). Reads below 1000 bp were filtered out and the best 5 Gbp of data with emphasis on length were kept, using Filtlong v0.2.1^27^. The initial assembly was de-novo assembled from the filtered long-reads using Flye v2.9.1 in nano-hq mode^28^. The assembly was polished in two rounds with the long-reads using Medaka v1.7.2 (Oxford Nanopore). Contigs with a coverage below 50x and contigs of bacterial origin, identified with blastx v2.14.0^29^ to the Swiss-Prot database (release-2022_05)^30^, were discarded. The assembly was polished with NextPolish^31^ using both short- and long-reads with “task=best”, which runs a total of two rounds of polishing with long-reads, followed by four rounds of polishing with short-reads in two different algorithm modes. The mean long-read coverage over all positions on the 12 *D. firmibasis* contigs was 159x, the mean short-read coverage was 441x, for a total of 599x coverage. NextPolish polishing resulted in 42,885 edits on the 31.5 Mbp assembly (Fig. 1).

The new assembly was compared to the old *D. firmibasis* ASM27748v1 assembly (GenBank GCA_000277485.1)^17^ using satsuma2 v2016-12-07^32,33^. Synteny analysis revealed that, although the old assembly is highly fragmented, there are no major regions of the new assembly entirely missing in the old assembly (Fig. 2). The two assemblies are also similar in size. The main difference between the assemblies is that the new assembly contains no undetermined bases (versus 4.1 Mbp undetermined in the old genome) and the new genome is much more contiguous, resulting in an assembly more representative of the *D. firmibasis* genome (Table 1).

**Table 1.** Old and new *D. firmibasis* assembly statistics.

|  | Old | New |
| --- | --- | --- |
| Genome size (Mbp) | 30.6 | 31.5 |
| Determined bases (Mbp) | 26.4 | 31.5 |
| Undetermined bases | 4,112,630 | 0 |
| # Gaps > 10bp | 6312 | 0 |
| # Contigs | 997 | 12 |
| N50 (Mbp) | 0.18 | 4.39 |
| A + T% | 75.2 | 76.0 |
| BUSCO complete% | 82.8 | 92.9 |

**Fig. 2.**
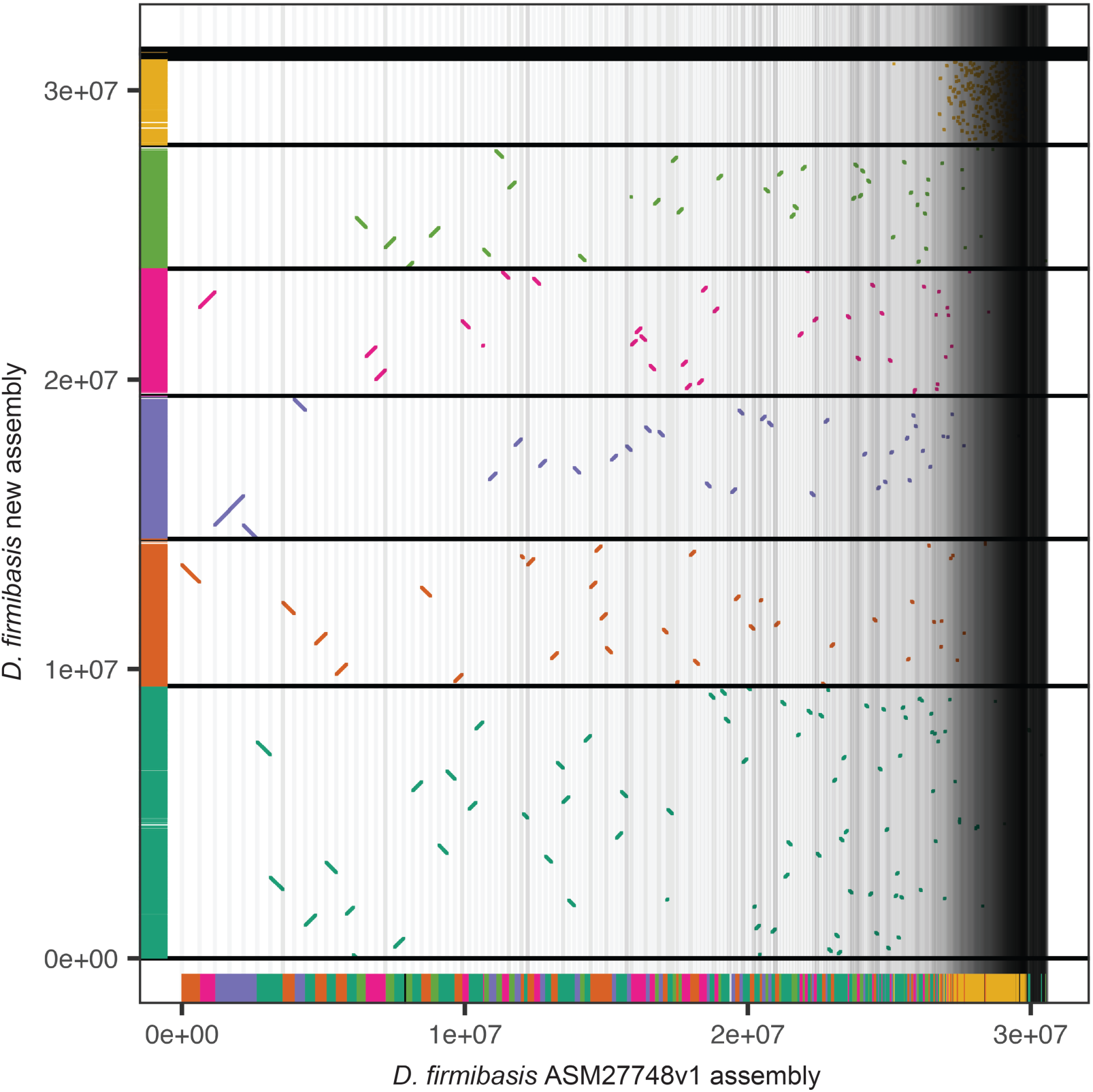
Comparison of the old and new *D. firmibasis* assembly. Regions of the *D. firmibasis* ASM27748v1 assembly^17^ with homology to the new assembly plotted, with a distinct color for each of the large contigs in the new assembly. Matches are plotted in 2D-space and on the x- and y-axes (old and new assembly respectively). Horizontal black lines indicate contig boundaries of the new assembly, vertical grey lines indicate contig boundaries of the *D. firmibasis* ASM27748v1 assembly.

### Genome annotation

For protein coding gene annotation, mRNA-seq reads were mapped to the genome using STAR v2.7.10b^34^. Mapped mRNA reads were assembled to transcripts using genome-guided Trinity v2.14.0^35^, with a maximum intron size of 5000bp. Annotation of the assembly was performed using MAKER v3.01.04^36^ with assembled transcripts and the *D. discoideum* UP000002195 UniProt reference proteome^30^ as gene evidence. Homology of annotated *D. firmibasis* proteins to proteins in the *D. discoideum* proteome was identified with blastp v2.14.0^29^.

In total, 379 tRNAs were annotated with tRNAscan-SE v2.0.12^37^, which in line with the number of tRNAs annotated in *D. discoideum* (Table 2). miRNAs were annotated with ShortStack v4.0.1^38^ based on the mature miRNAs reported previously^19^, and *D. firmibasis* small RNA-seq reads from NCBI BioProject PRJNA972620^19^. Class I RNAs were annotated based on sequence, structure and presence of up-stream potential promotor motifs as described previously^10^ (Table 2). Other non-coding RNAs, such as rRNAs, snRNAs and snoRNAs, were annotated using infernal v1.1.4^39^ with a covariance model from Rfam^40^ filtered to Amoebozoa. Transposable elements were identified with tblastx v2.14.0^29^ with e-values below 10^−15^ and reference transposable element sequences from Repbase (Genetic Information Research Institute) filtered to *D. discoideum*.

**Table 2.** Number of annotations in *D. firmibasis* and *D. discoideum D. firmibasis D. discoideum*.

|  | *D. firmibasis* | *D. discoideum* |
| --- | --- | --- |
| Protein coding genes | 10,564 | 11,326 (dictyBase curated<sup>25</sup>) |
| tRNAs | 379 | 418<sup>16</sup> |
| miRNAs | 9 | 28<sup>11,12,14,46</sup> |
| Class I RNAs | 10 | 38<sup>13</sup> |
| snRNAs | 28 | 17<sup>15,47</sup> |
| snoRNAs | 23 | 47<sup>15,48</sup> |

The combined number of protein coding genes and (nc)RNA genes amounted to a total number of 11077 annotated genes. Gene annotations were manually verified and adjusted where necessary, using NCBI Genome Workbench v3.8.2^41^. The annotated genome has been submitted to NCBI with genome accession number JAVFKY000000000^42^. The combined annotation resulted in a mean gene density of 70.6% which is distributed homogeneously over the main contigs, but reduced towards the ends of the contigs (Fig. 3a). Additionally, there is a region with low gene density on chr1 between position 4.5 Mbp and 4.8 Mbp, which is matched with lower mRNA-seq coverage. This region contains a large DIRS1 retrotransposon (Fig. 3a). DIRS1 is known to be targeted by small interfering RNAs in *D. discoideum*^14,43^. Small RNA data from *D. firmibasis*^19^ were mapped to the genome using ShortStack v4.0.1^38^. Coverage of small RNAs was substantially higher on the region featuring DIRS1 than the surrounding regions. This is in agreement with the situation in *D. discoideum* and could be partly responsible for the low mRNA-seq coverage in the region (Fig. 3b). Other large DIRS1 insertions were found at the ends of chromosomes 2, 3, 4, and 5 (Fig. 3a). This is similar to what has been reported for *D. discoideum*, which contains clusters of DIRS1 repeats near one end of each chromosome^16^. When the *D. discoideum* genome was sequenced, no conventional telomeric repeats could be found at the ends of the six chromosomes^16^. In the new *D. firmibasis* assembly however, potential telomeric repeats with sequences 5’-GAGGAGAGAGTCCCTTTTTTT-3’ and 5’-GGGGAGAGACA-3’ could be identified. The repeats were annotated using bowtie v1.3.1^44^ allowing two mismatches in repeat 5’-GAGGAGAGAGTCCCTTTTTTT-3’ and one mismatch per 5’-GGGGAGAGACAGGGGAGAGACA −3’ double repeat. Telomeric repeats were found at both ends of chromosomes 1 and 6, and one end of chromosomes 3 and 4 (Fig. 3a), and were not present elsewhere on the chromosomes. Both types of telomeric repeats could be identified on the same chromosome end, and an average of 15 repeats could be identified per end.

**Fig. 3.**
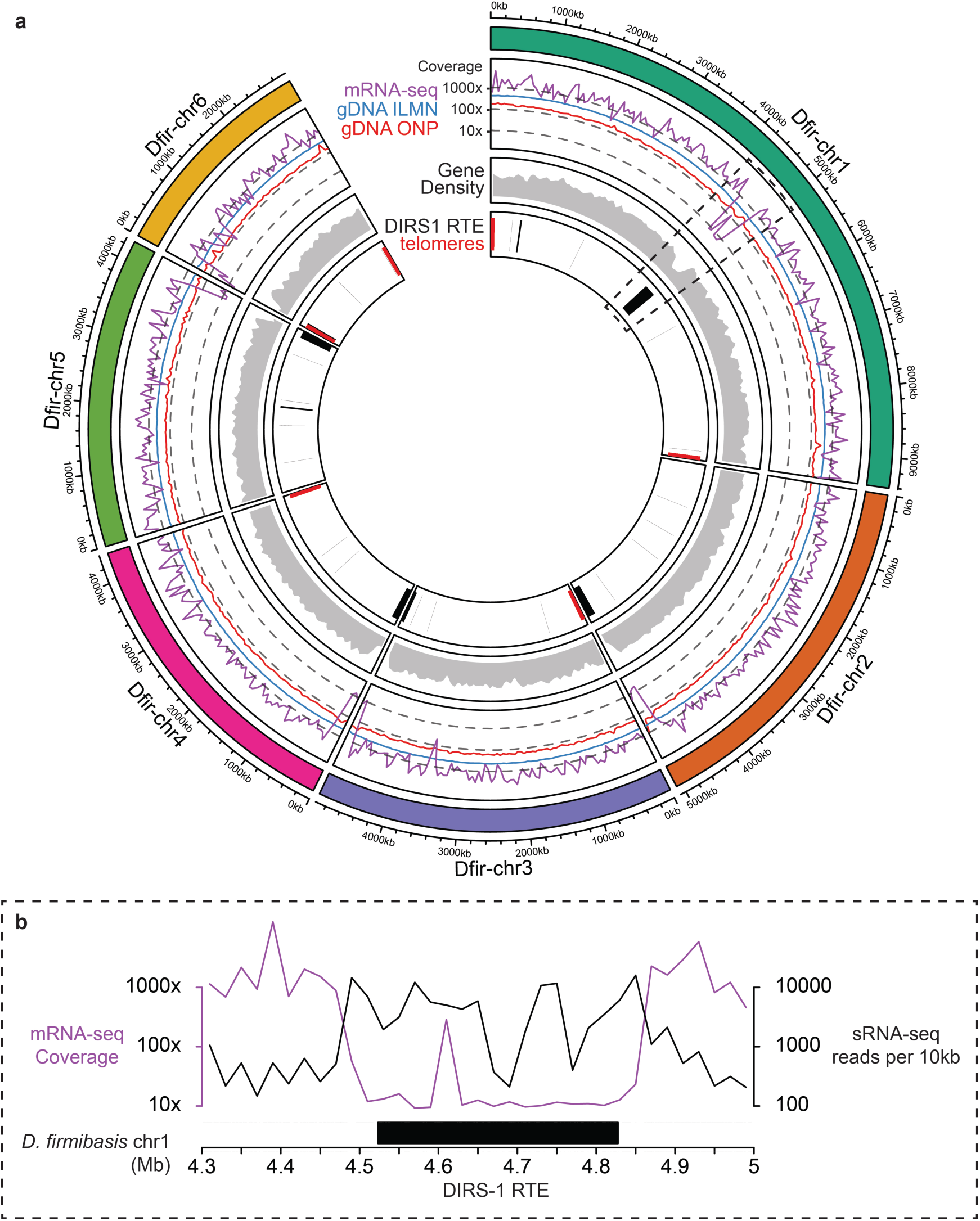
Coverage and annotation of the new *D. firmibasis* assembly. **a** Circular representation with the six chromosomal contigs (chr) in the new *D. firmibasis* assembly. From outside to inside: coverage of mRNA-seq (purple line), gDNA from Illumina short-read sequencing (gDNA ILMN, blue line) and gDNA from Oxford Nanopore long-read sequencing gDNA (gDNA ONP, red line); gene density (grey); annotation of DIRS1 retrotransposable elements (DIRS1 RTE, black rectangles) and telomeric repeats (red rectangles). Circular plot generated with the R package *circlize*^45^. **b** Close-up of the *D. firmibasis* chr1 contig from position 4.3 Mbp to 5 Mbp, containing a DIRS1 retrotransposable element (DIRS1 RTE, solid black bar), with coverage from mRNA-seq (purple line) and number of small RNA-seq (black line, sRNA-seq) reads mapping per 10kb region. sRNA-seq reads accessed with NCBI Bioproject accession PRJNA972620^19^.

## Data Records

This Whole Genome Shotgun project has been deposited at DDBJ/ENA/GenBank under the accession JAVFKY000000000^42^. The version described in this paper is version JAVFKY010000000. The gDNA and mRNA sequencing libraries have been submitted to SRA with BioProject accession number PRJNA1008051^49^. In total, 18 paired-end mRNA sequencing libraries are associated with this work – 9 for *D. firmibasis* and 9 for *D. discoideum* – from three distinct morphological stages: vegetative, slug and fruiting body stages, identifiable with respectively “0h”, “16h” and “24h” in the repository.

## Technical Validation

### Chromosome-level contiguity and completeness

By comparing the new assembly presented here to the first published *D. firmibasis* assembly^17^, we could show that no major parts of the old assembly were missing in the new assembly (Fig. 2). Similarly, a synteny search was performed between the new *D. firmibasis* assembly and the *D. discoideum* AX4 genome^16,50^. *D. discoideum* is relatively closely related to *D. firmibasis*^18^, and in line with that we could identify extensive synteny between the two species (Fig. 4a). There appears to have been no major reorganization of *D. firmibasis* chromosomes 2, 3 and 5, which match *D. discoideum* chromosomes 4, 5 and 1, respectively (Fig. 4a). *D. firmibasis* chromosome 1 matches *D. discoideum* chromosomes 3 and 6, with large inversions. The region of *D. firmibasis* chromosome 1 that harbors the DIRS1 retrotransposon appears to be missing in *D. discoideum*, revealing that this insertion might have occurred more recently or was lost in *D. discoideum*. Chromosome 2 of *D. discoideum* features a 1.5 Mbp inverted duplication^16^. This can be observed here, since a region of the *D. firmibasis* chromosome 6 is represented twice in *D. discoideum* chromosome 2 (Fig. 4a). Strikingly, it is in this same region that the *D. firmibasis* chromosomes 4 and 6 are split, relative to the larger *D. discoideum* chromosome 2 (Fig. 4a). This is in line with the hypothesis, that the region where the duplication is found, is prone to breakage^16^. Since all *D. discoideum* chromosomes are accounted for in the six large *D. firmibasis* contigs, we conclude the new *D. firmibasis* assembly is complete, and of chromosome-level quality.

**Fig. 4.**
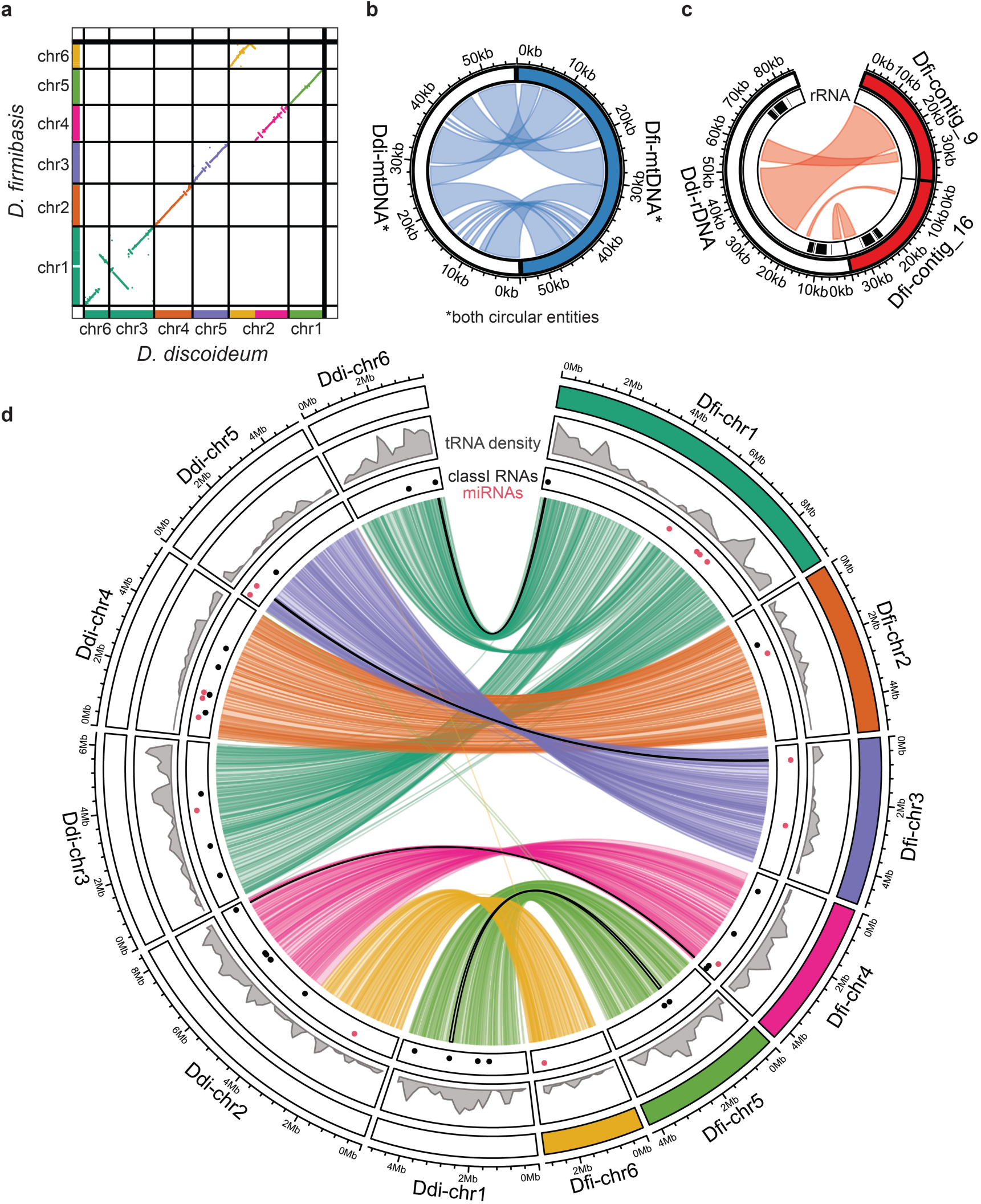
Extensive conservation of synteny between *D. discoideum* AX4 and *D. firmibasis*. **a** Comparison of genomes as in Fig. 2 but between the new *D. firmibasis* assembly and the dicty_2.7 *D. discoideum* AX4 assembly, with chromosome numbers indicated. The *D. firmibasis* chromosomes are colored as previously. Regions of the *D. discoideum* dicty_2.7 assembly are colored to match the *D. firmibasis* chromosomes they are homologous to. **b** Circular alignment of the mitochondrial genomes from the *D. discoideum* (Ddi-) and *D. firmibasis* (Dfi-) assemblies. Homologous regions are visualized with light blue links. Regions within 500 bp proximity were merged for visual clarity. **c** As in **b**, but for the *D. discoideum* extrachromosomal DNA palindrome containing the ribosomal DNA (rDNA), aligned to *D. firmibasis* contig_9 and contig_16. rDNA locations are represented by black lines in the inner ring. Regions of homology (light red) within 5 kbp proximity in the same direction on both contigs were merged. **d** Circular alignment of the chromosomes of the dicty_2.7 *D. discoideum* AX4 assembly and the new *D. firmibasis* assembly. From outside to inside is the visual representation of the contigs, colored for *D. firmibasis*, white for *D. discoideum*; density of tRNAs on the genome assemblies (grey); presence of Class I RNAs (black) and miRNAs (red); and links between the two assemblies to indicate synteny regions based on nucleotide homology. The links are colored according to the *D. firmibasis* contig it matches. Four of the synteny regions contain miRNAs in both assemblies or Class I RNAs in both assemblies – outlined black. Regions of homology within 5 kbp proximity in the same direction on both contigs were merged. Circular plots were generated with the R package *circlize*^45^.

Besides the chromosomes, synteny in four of the six remaining small contigs was detected as well. The *D. firmibasis* mitochondrial DNA was assembled as a circular entity, indicating that the entire mitochondrial genome was covered. Between *D. firmibasis* and *D. discoideum* there appear to have been no major reorganizations in the mitochondrial genomes (Fig. 4b). Furthermore, the linear extrachromosomal DNA containing the rRNA genes in *D. discoideum* matched two smaller *D. firmibasis* contigs, which also contain annotations for the 5S, 5.8S, 18S and 28S rRNA genes (Fig. 4c). In *D. discoideum*, this extrachromosomal DNA is palindromic, with rRNA genes on both sides, but in the combined *D. firmibasis* contigs, only one set of the rRNA genes is assembled.

Much of the homology between the two genomes was detected due to conservation of protein coding regions. However, we were also interested in understanding to what extent the genes coding for ncRNAs were conserved between the two species. Total number of tRNAs is high in both species, with 379 and 418 annotated tRNA genes in the *D. firmibasis* and *D. discoideum* genome^16^, respectively (Table 2). Not only do their numbers match, they also appear to be located in homologous areas of the genome, as seen from the tRNA density plot (Fig. 4d). Small ncRNAs such as Class I RNAs and miRNAs appear to be much less conserved and rapidly evolving, as previously reported^10,19^ (Table 2). Here, we could only detect four synteny regions which contained Class I RNAs or miRNAs in both species (outlined with black lines connecting the genomes in Fig. 4d). Besides these, the majority of Class I RNAs and miRNAs appear to be unique to each species.

### Validation of gene annotation and expression

*D. firmibasis* and *D. discoideum* mRNA-seq reads from three distinct morphological stages (Fig. 1; three biological replicates from each stage) were mapped to their respective genomes using STAR v2.7.10b^34^. Reads were assigned to genes with featureCounts v2.0.3^51^. Using the new *D. firmibasis* assembly, 95% of all reads could be mapped to the genome, and 93% of all mapped reads were assigned to genes, demonstrating completeness of the genome and annotations.

Of the 11077 annotated *D. firmibasis* genes, homology evidence to *D. discoideum* genes could be identified with blastp v2.14.0^29^ for 10192 genes. Some *D. firmibasis* genes matched the same *D. discoideum* gene, in which case the counts were summed. Additionally, not all *D. firmibasis* or *D. discoideum* genes were expressed, resulting in 8359 homologs to be compared. Logarithmic fold change (logFC) of genes at the slug stage and fruiting body stage relative to the vegetative stage was calculated using DESeq2 v1.42.0^52^ and scaled using *apeglm*^53^. Expression of the 8359 homologous genes was compared between *D. discoideum* and *D. firmibasis* (Fig. 5a). These genes appear to be expressed in a broadly similar fashion during multicellular development, showing that the regulation of genes that govern the transition from unicellular to multicellular in *D. firmibasis* and *D. discoideum* is largely the same (Fig. 5a). Of the 8359 homologs, 202 genes showed significantly different regulation (adjusted p-value < 0.01) (Fig. 5b). The regulatory pattern of these genes during development may have diverged between these species, and are interesting candidates for further studies.

**Fig. 5.**
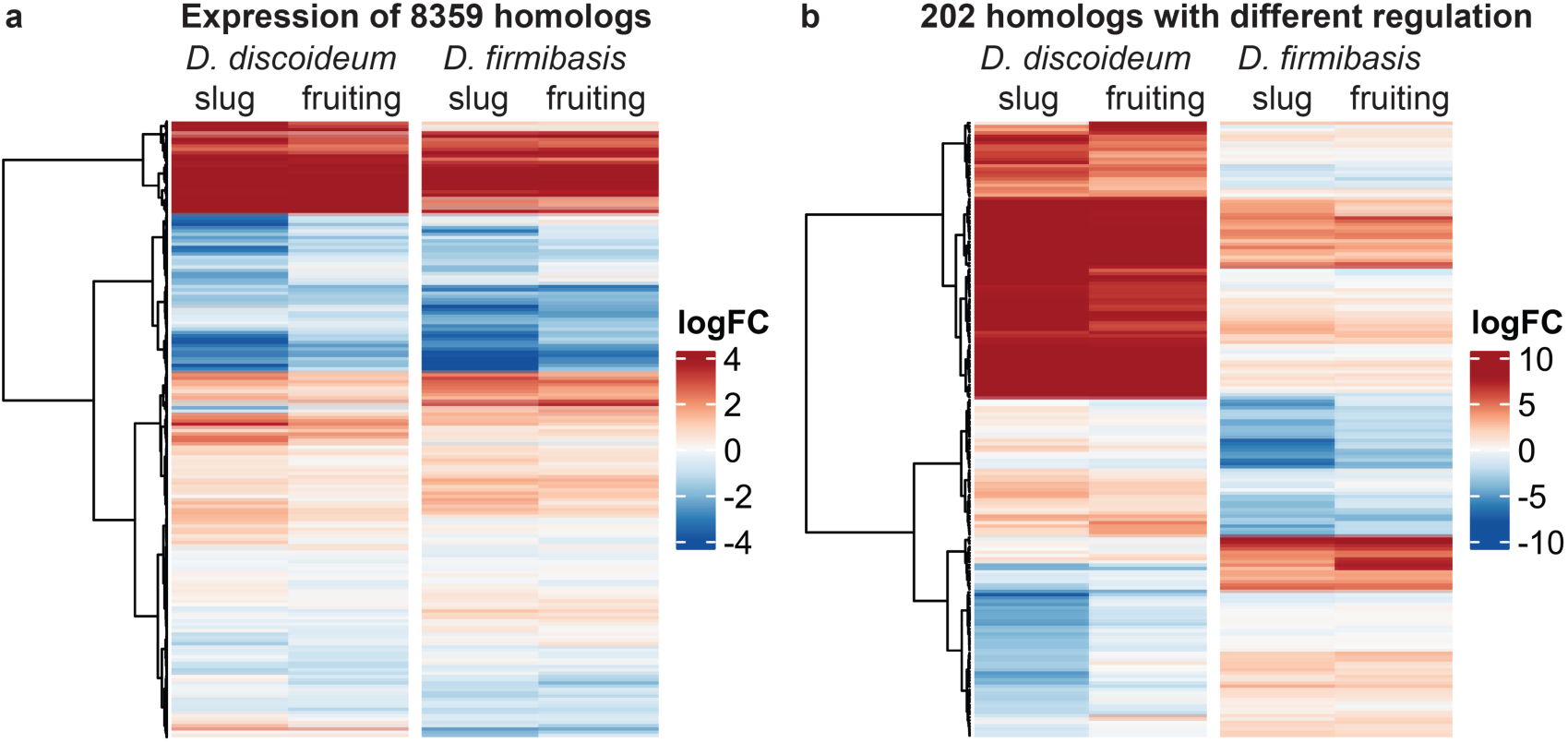
Comparison of the developmental transcriptome between *D. discoideum* and *D. firmibasis*. Expression of homologs in both species at the slug stage (slug) and in fruiting bodies (fruiting) versus their respective vegetative stages, hierarchically clustered based on their logFC. **a** Expression of all 8359 homologs expressed in both species. **b** Expression of 202 homologs with a significantly different regulation during development (adjusted p-value < 0.01).

## Code Availability

Software was run with standard parameters unless otherwise indicated. Custom code and scripts, to analyze the genome, annotations, or generate figures is available at figshare^54^.

## Acknowledgements

Sequencing was performed by the SNP&SEQ Technology Platform in Uppsala. The facility is part of the National Genomics Infrastructure (NGI) Sweden and Science for Life Laboratory. The SNP&SEQ Platform is also supported by the Swedish Research Council and the Knut and Alice Wallenberg Foundation. The computations and data handling were enabled by resources provided by the National Academic Infrastructure for Supercomputing in Sweden (NAISS) at Uppsala University, partially funded by the Swedish Research Council through grant agreement no. 2022-06725. Additionally, we would like to acknowledge DictyBase (http://dictybase.org/) for providing strains. This work was supported by grant no. 2021-05793 from Swedish Research Council (Vetenskapsrådet) to Fredrik Söderbom.

## Author contributions

B.E.: extraction of DNA and RNA from *D. firmibasis*, long-read sequencing and library preparation, genome annotation and analysis, preparation of the figures and drafting of the manuscript. J.K: long-read sequencing and library preparation, and genome annotation. J.J-H: long-read sequencing, assembly, acquired funding. S.K.: supervision, acquired funding. F.S.: supervision, acquired funding. B.E., J.K., J.J-H, and F.S. were involved in the study design and all authors have revised and approved the final manuscript.

